# Fluorescent tagging of endogenous IRS2 with an auxin-dependent degron to assess dynamic intracellular localization and function

**DOI:** 10.1101/2023.12.06.570406

**Authors:** Minjeong Jo, Ji-Sun Lee, Michael W. Lero, Jennifer S. Morgan, Leslie M. Shaw

**Author notes:** Address Correspondence to: Leslie M. Shaw, PhD Department of Molecular, Cell & Cancer Biology University of Massachusetts Chan Medical School 364 Plantation St., LRB 409 Worcester, MA 01605 Voice.

## Abstract

Insulin Receptor Substrate 2 (IRS2) is a signaling adaptor protein for the insulin (IR) and Insulin-like Growth Factor-1 (IGF-1R) receptors. In breast cancer, IRS2 contributes to both initiation of primary tumor growth and establishment of secondary metastases through regulation of cancer stem cell (CSC) function and invasion. However, how IRS2 mediates its diverse functions is not well understood. We used CRISPR/Cas9-mediated gene editing to modify endogenous IRS2 to study the expression, localization, and function of this adaptor protein. A cassette containing an auxin inducible degradation (AID) sequence, 3X-FLAG tag and mNeon-green was introduced at the N-terminus of the IRS2 gene to provide rapid and reversible control of IRS2 protein degradation and analysis of endogenous IRS2 expression and localization. Live fluorescence imaging of these cells revealed that IRS2 shuttles between the cytoplasm and nucleus in response to growth regulatory signals, and deletion of a putative nuclear export sequence in the C-terminal tail promotes nuclear retention of IRS2. Moreover, acute induction of IRS2 degradation reduces CSC function, similar to the constitutive knockout of IRS2. Our data highlight the value of our model of endogenously tagged IRS2 as a tool to elucidate IRS2 localization and function.

## Introduction

The Insulin Receptor Substrate (IRS) adaptor proteins IRS-1 and IRS-2 play essential roles in regulating the response of breast and other tumor cells to signaling through the insulin (IR) and Insulin-like Growth Factor-1 (IGF-1R) receptors, as well as to some integrin and cytokine receptors (1,2). Both IRS1 and IRS2 are expressed in breast tumors, but their expression patterns and functions differ in a subtype-dependent manner. IRS1 expression is highest in well-differentiated ER+ luminal breast tumors, and expression decreases as tumors become more poorly differentiated (3,4). IRS1 has been implicated in regulating proliferation and survival in response to insulin and IGF-1 signaling (IIS) (5-7). In contrast, IRS2 expression is elevated in more aggressive subtypes such as Basal/Triple Negative Breast Cancer (TNBC) and the localized expression of IRS2 is associated with reduced overall survival of breast cancer patients (8,9). Irs2 contributes to both initiation of primary tumor growth and establishment of secondary metastases through regulation of CSC function and invasion in response to IIS (10-13). In contrast, loss of Irs1 expression enhances metastatic rates (14). Together, studies to-date highlight the divergent functions of the IRS adaptor proteins in breast cancer. However, how each adaptor protein regulates diverse functions is not well understood and more detailed studies are required to address this lack of mechanistic insight.

Much of the understanding about how the IRS proteins impact tumor cell functions has come from studies of cell lines in which IRS expression has been stably knocked down by shRNA or knocked out by CRISPR/Cas9 mediated gene targeting and restoration of exogenously expressed WT and mutant proteins (11-13). These studies have provided important information about these proteins, but a rigorous understanding of endogenous protein localization in living cells and the functional impact of acute loss of protein expression is lacking. CRISPR-mediated endogenous gene tagging provides the potential to explore protein function and subcellular localization in living cells or more complex model systems. For example, fluorescently tagged proteins are detectable in live cells enabling a complete investigation of their subcellular location (15). In addition, the auxin-inducible degron (AID) system has provided a unique tool for rapid and reversible depletion of a target protein, controlled by the addition of an auxin molecule (indole-3-acetic acid, 3-IAA) (16,17). When an AID is fused to a target protein and an auxin receptor F-box protein is exogenously expressed, a SKP1-CUL1-F-Box (SCF) ubiquitin E3 ligase complex is formed in eukaryotic cells using endogenous components. Following the addition of IAA, polyubiquitination and rapid proteasomal degradation of the degron-fused protein occurs within just a few hours. Acute loss-of-function experiments reveal significant insights into molecular pathways and gene function, and the AID system has been successfully applied in many cell lines and organisms (18).

In this study, we utilized CRISPR/Cas9-mediated homology-directed repair to generate cells expressing endogenous IRS2 containing an N-terminal auxin-inducible degron and mNeon-green fluorescent protein to monitor IRS2 expression and function. The resources developed in this study should be of value in studying the complexity of IRS2 function and dynamic regulation.

## Results

### CRISPR/Cas9-mediated targeting of an AID-FLAG-mNeon-green tag at the IRS2 N-terminus

To facilitate real-time analysis of endogenous IRS2 expression and localization and study the impact of acute loss of IRS2 expression on cellular function, we utilized CRISPR/Cas9-mediated homology directed repair (HDR) to insert a cassette that included an auxin-inducible **D**egron (AID) sequence from *Arabidopsis thaliana* IAA17 (mIAA7 degron), a 3X**F**LAG tag and the fluorescent m**N**eon-green protein (DFNcassette) at the N-terminus of the IRS2 protein (Fig. 1A). A flexible linker was added between this cassette and the N-terminus of IRS2 to prevent interference with IRS2 function. Guide RNAs that recognized protospacer adjacent motif (PAM) sequences within 20 bp of the start codon of IRS2 were screened using Tracking of Indels by Decomposition (TIDE) analysis to identify the guide with the most efficient cutting (Supplemental Fig 1A).

**Figure 1.**
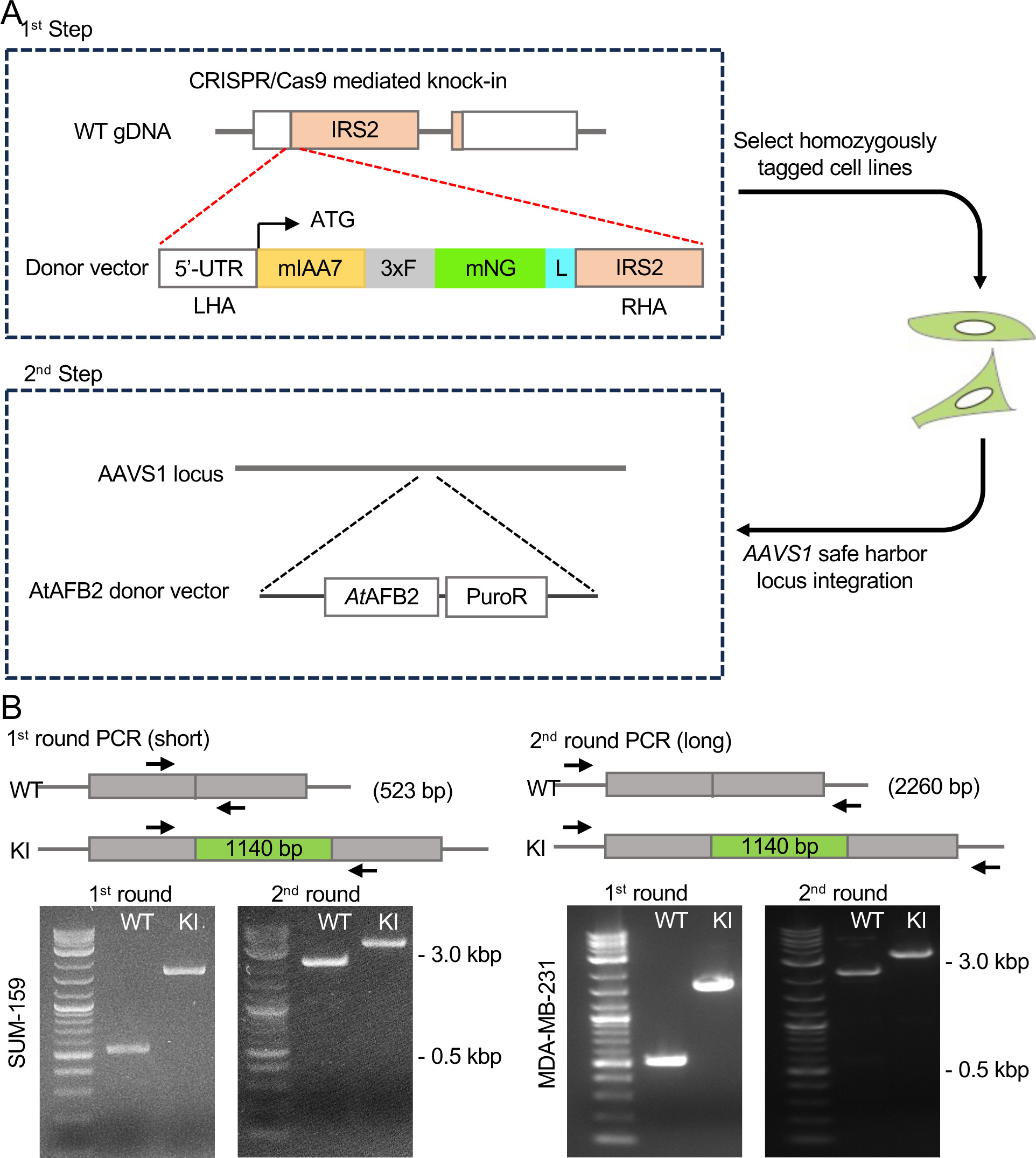
Schematic of the generation of IRS2 knock-in cells. A. Targeted integration of AID, FLAG, and mNeonGreen tags with linker into the N-terminus of IRS2 and integration of AtAFB2 and the PuroR cassette into the AAVS1 safe harbor locus in homozygously tagged clones by selection with puromycin. B. PCR genotyping strategy for selecting homozygously tagged *IRS2*. 1^st^ round primers complementary to the modified loci were used to amplify wild type alleles (523 bp) and knock-in alleles (1663 bp). 2^nd^ round primers complementary to IRS2 outside the donor plasmid homology arms were used in long-distance PCR for wild type alleles (2260 bp) and knock-in alleles (3400 bp). Indicated amplicon sizes confirm the homozygous integration of the tags.

Knock-in cells were generated in two steps using CRISPR/Cas9-mediated HDR. In the first step, SUM-159 and MDA-MB-231 triple negative breast carcinoma cells were electroporated with the most efficient gRNA complexed with Cas9 protein, and a donor fragment containing the DFN-cassette with left and right homology regions corresponding to the IRS2 genomic sequence (Fig 1A). After electroporation, cells were plated and allowed to recover for 24-48 hrs before sorting mNeon-green positive cells into 96 well plates (Supplemental Fig. 1B). Expanded clones were screened by genomic PCR to confirm knock-in of the DFN cassette into the N-terminus of the IRS2 protein. In the first round of PCR screening, primers amplified the DFN cassette alone and in the second round, primers amplified the DFN cassette and both homology arms (Fig. 1B). A single band with an increased shift of 1140 bp, the size of the DFN cassette, in the knock-in cells indicates homozygous expression of tagged-IRS2, which was confirmed by Sanger sequencing. In the second step, selected single homozygously-tagged clones were modified to stably express the *Arabidopsis thaliana* F-box protein AFB2 (*At*AFB2), which upon treatment with auxin, binds to the mIAA7 degron and promotes ubiquitination and degradation of tagged proteins (19). AtAFB2 was inserted into the AAVS1 safe harbor locus by CRISPR/Cas9-mediated HDR (Fig. 1A) (17,19). After selection in puromycin, cells were treated with Auxin for 1 hour and then sorted for loss of mNeon-green fluorescence to further enrich for AtAFB2 positive populations.

### Characterization of cells expressing endogenously tagged IRS2

Cell extracts were screened by immunoblotting to confirm the expression of the DFN-IRS2 tagged protein. Insertion of the DFN cassette resulted in the expected mobility shift of 42 kDa for the IRS2 protein, which was also recognized by FLAG and mNeon-green specific antibodies (Fig. 2A). mNeon-green expression in the knock-in cells was also confirmed by fluorescence imaging and flow cytometry (Figs 2B and 2C). WT parental cells were negative for mNeon-green and FLAG expression by both immunoblot and imaging.

**Figure 2.**
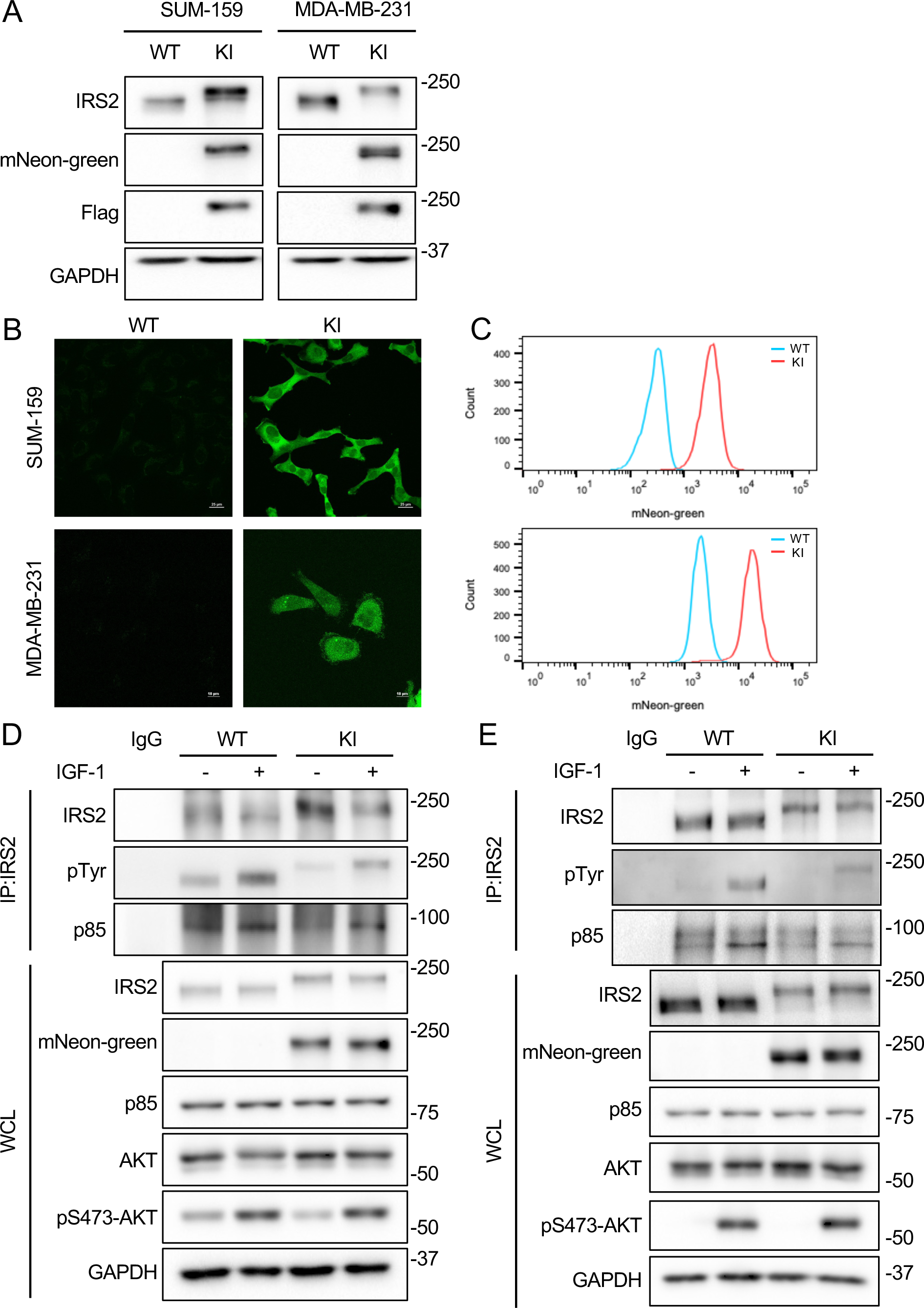
Characterization of DFN-IRS2 protein expression and function. A. WT and DFN (KI) SUM-159 and MDA-MB-231 cells were immunoblotted with antibodies that detect IRS2 mNeon-green and FLAG. B. Live fluorescence imaging of WT and DFN (KI) SUM-159 and MDA-MB-231 cells. C. Flow cytometric analysis of WT and DFN (KI) SUM-159 and MDA-MB-231 cells. D and E. WT and DFN (KI) SUM-159 and MDA-MB-231 cells were serum starved and stimulated with IGF-1 (50 ng/ml). Aliquots of cell extracts that contained equivalent amounts of total protein were immunoprecipitated with IRS2-specific antibodies and the immune complexes were immunoblotted with antibodies that recognize either IRS2, phosphotyrosine (pTyr) or p85. Aliquots of cell extracts that contained equivalent amounts of total protein (WCL) were immunoblotted with the indicated antibodies.

IRS2 is recruited to the phosphorylated cytoplasmic tails of the insulin and IGF-1 receptors upon stimulation with insulin/IGF-1 (20). Upon binding, IRS2 undergoes tyrosine phosphorylation by the intrinsic receptor tyrosine kinases, creating new docking sites for downstream signaling effectors such as PI3K (21). To confirm that addition of the DFN cassette to the N-terminus of IRS2 does not interfere with this signaling function, cells expressing WT-IRS2 and DFN-IRS2 were stimulated with IGF-1 (50 ng/ml) and cell extracts were immunoprecipitated with IRS2 antibodies. Immune complexes were resolved by SDS-PAGE and immunoblotted with phosphotryosine (pTyr) and PI3K (p85)-specific antibodies. Increased phosphorylation and binding to PI3K was observed after stimulation with IGF-1 for WT-IRS2 and DFN-IRS2 in both SUM-159 and MDA-MB-231 cells (Figs 2D and 2E). Activation of AKT downstream of PI3K, as measured by phosphorylation of S473-AKT, was also observed for both KI cell lines.

### Reversible IRS2 protein degradation by auxin

Tagging IRS2 with an auxin-dependent degron allows for inducible and reversible suppression of endogenous IRS2 expression to study the impact of acute IRS2 loss on cellular function. To assess the efficiency of IRS2 degradation and recovery, DFN-IRS2 cells were treated with the auxin derivative 3-IAA (500 μM). 3-IAA treatment induced the degradation of IRS2 in both SUM-159-DFN and MDA-MB-231-DFN cells in a time dependent manner, with IRS2 levels significantly diminished after 6 hours and reaching a plateau by 24 hours (Fig. 3A and Supplemental Fig 1C). Upon wash-out of 3-IAA, IRS2 expression recovered in a time-dependent manner within 6 hrs (SUM-159) or 24 hrs (MDA-MB-231) (Fig 3B). The recovery was faster in SUM-159 cells when compared with MDA-MB-231 cells, which may reflect differences in rates of IRS2 transcription or translation in these cells. A low level of IRS2 expression remained in both cell lines after auxin treatment, which may reflect the inability of the F-box protein to access the AID tag on IRS2 due to either structural interference or spatial restriction of the two proteins.

**Figure 3.**
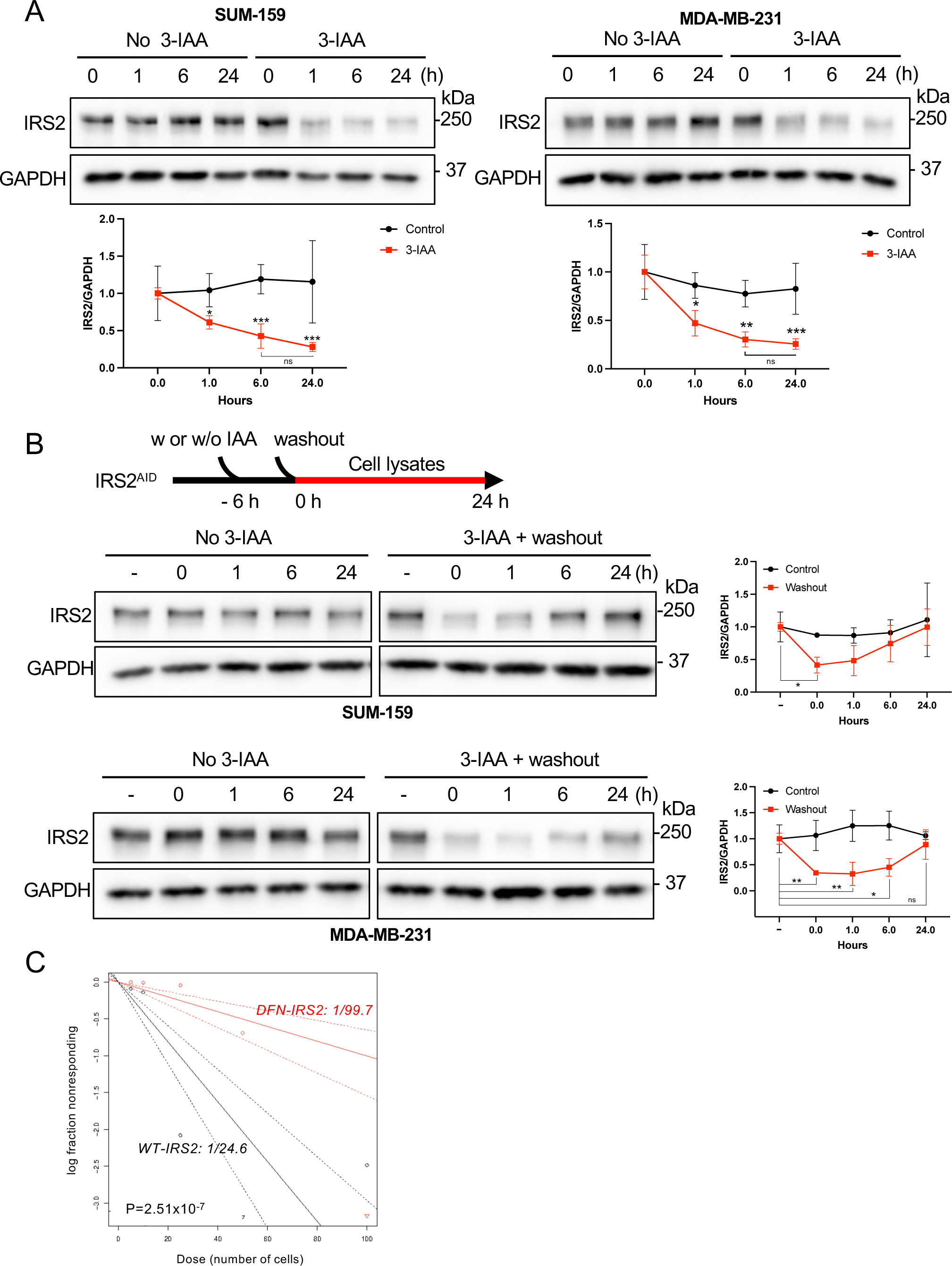
Auxin-mediated degradation of DFN-IRS2. A. DFN KI-cells were treated with or without IAA (0.5 mM) for the indicated time periods and cell extracts were immnunoblotted with IRS2 antibodies. The data shown in the graphs represent the mean ± S.D. of three independent experiments. B. DFN KI-cells were treated with or without 0.5 mM IAA for 6 hrs. The cells were then washed and allowed to recover for the indicated time periods in complete culture medium. The data shown in the graphs represent the mean ± S.D. of three independent experiments. C. WT and DFN-KI SUM-159 cells were treated with 0.5 mM IAA for 24 hrs and then analyzed by *in vivo* limiting dilution assay. *p < 0.05, **p < 0.01 ***p < 0.001

A role for IRS2 in cancer stem cell (CSC) regulation was established in our previous studies using cells chronically lacking IRS2 expression through Cre-lox recombination or CRISPR/Cas9-mediated gene knockout (12). To evaluate the impact of acute reduction of IRS2 expression on CSC function, *in vitro* limiting dilution assays were performed to quantify CSCs in the population after exposure to auxin to induce IRS2 degradation (12). Parental and DFN-IRS2 SUM-159 cells were treated with 3-IAA for 24 hrs prior to performing *in vitro* limiting dilution assays. As shown in Fig 3C, stem cell numbers were diminished 4-fold when IRS2 expression was acutely suppressed by auxin-mediated degradation in the IRS2-DFN cells (1/24.6 for WT-IRS2 expressing cells and 1/99.7 for DFN-IRS2 cells).

### Live imaging of IRS2 reveals dynamic intracellular localization

Knocking mNeon-green into the N-terminus of IRS2 allows for the analysis of endogenous IRS2 localization in live cells. This analysis eliminates the potential for artifacts from fixation and Ab immunostaining. We initially assessed IRS2 expression in live cells under normal growth conditions or after serum-starvation and restimulation with IGF-1. As we had observed previously in fixed cells and in FFPE sections of human tumors, in normal growth medium IRS2 is primarily cytoplasmic with little expression in the nucleus (Figs 2B and 4A,B) (9,22). In contrast, when cells were serum-starved overnight, the nuclear localization of IRS2 increased markedly (Fig 4A,B). Stimulation with IGF-1 resulted in a time-dependent decrease in nuclear expression of IRS2 and the return toward a predominantly cytoplasmic localization pattern. These observations suggest that the ability of IRS2 to shuttle between the cytoplasm and nucleus can be controlled by intracellular signaling.

**Figure 4.**
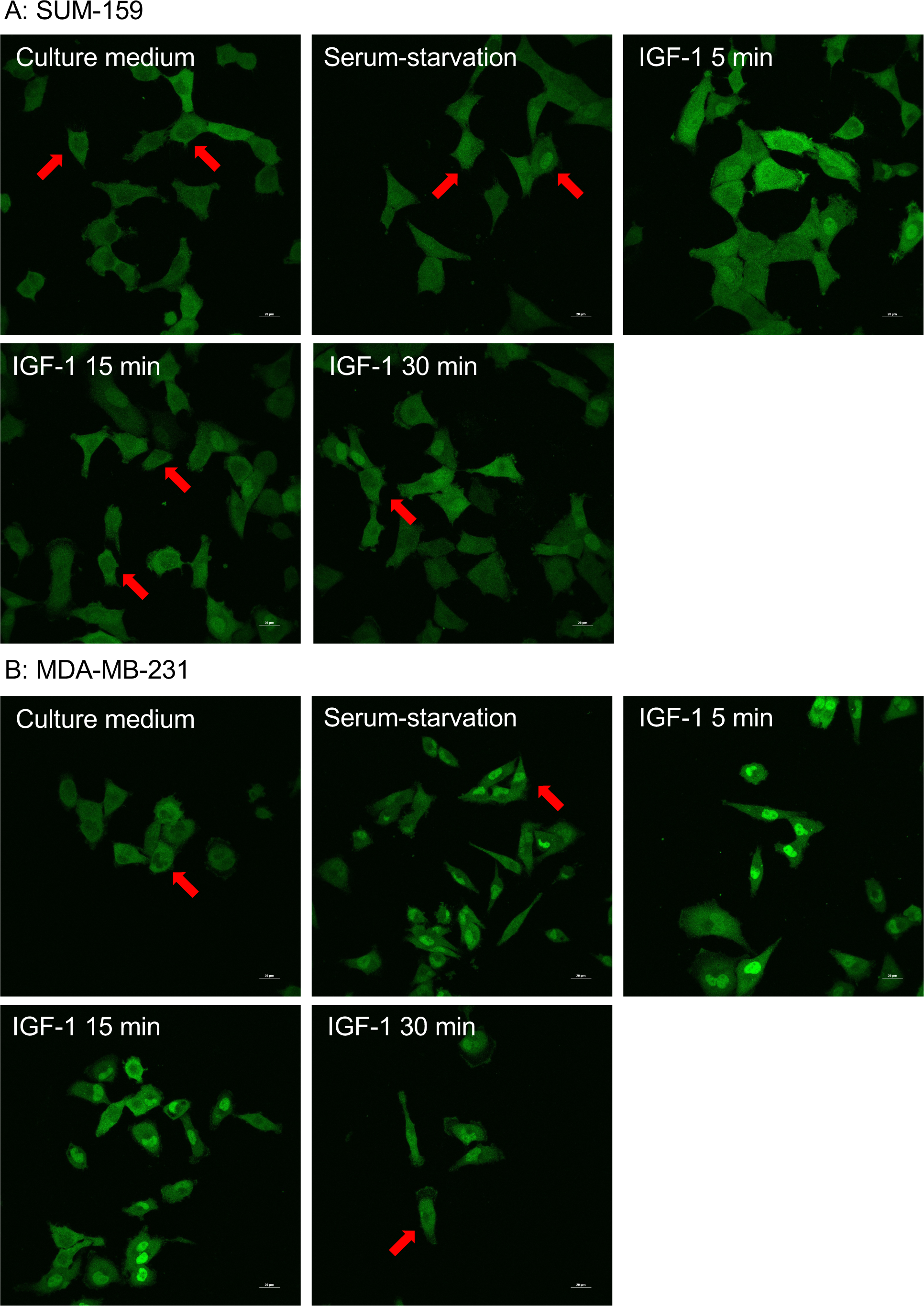
Live imaging analysis of DFN-KI cells. DFN-SUM-159 (A) and DFN-MDA-MB-231 (B) cells were analyzed by live fluorescence imaging in normal culture medium or culture medium lacking FBS (starvation) and then stimulated with IGF-1 (50 ng/ml). Red arrows, nucleus.

To understand how IRS2 nuclear-cytoplasmic localization is controlled, we analyzed the sequence of IRS2 using the LocNES prediction program to identify a putative CRM1-dependent nuclear localization signal (NES) (23). The sequences between 1096-1110 scored highly using this predictive program (Fig 5A). To assess if this region of IRS2 is required for nuclear export, we analyzed a series of C-terminal tail deletions of IRS2 (Fig 5B) for their localization in normal culture medium, which promotes cytoplasmic localization. IRS2 lacking the final 150 amino acids (ι11188) remained primarily in the cytoplasm, similar to WT-IRS2. However, deletion of an additional 174 amino acids (ι11015), which includes the putative NES, resulted in a significant increase in IRS2 expression within the nucleus, indicating that sequences in this region are required for efficient export of IRS2 from the nucleus.

**Figure 5.**
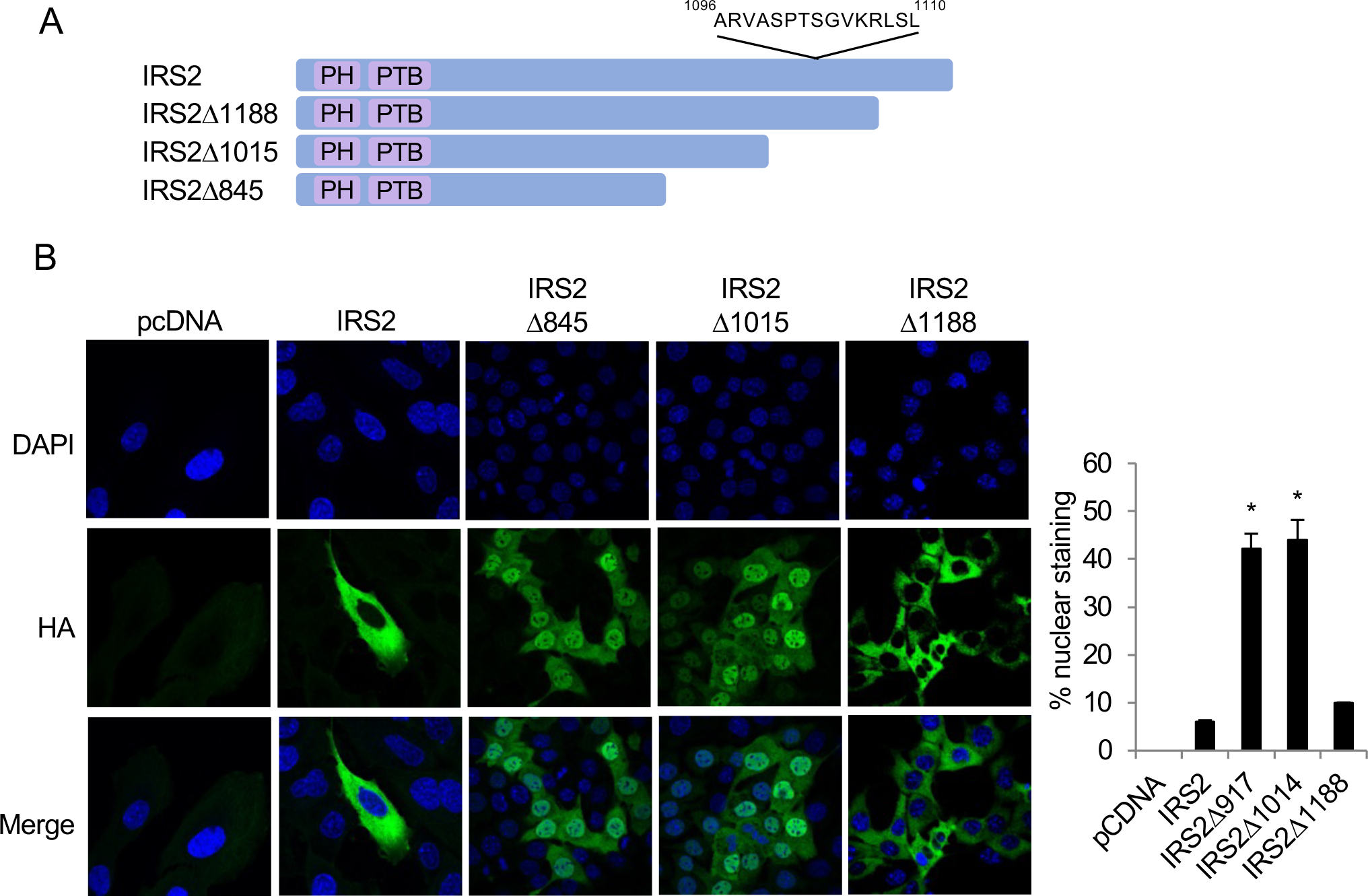
Intracellular localization of IRS2 and C-terminal mutants. A. Schematic of IRS2 and IRS2-deletion mutants showing the putative nuclear export signal. B. SUM-159 cells transfected with HA-tagged IRS2 and deletion mutants were fixed and stained with HA antibodies (green) and DAPI (blue). The percentage of IRS2 expression in the nucleus is shown in the graph (right). *p < 0.05

## Discussion

We used CRISPR/Cas9-mediated gene editing to modify endogenous IRS2 to study the expression, localization and function of this adaptor protein. Introduction of an auxin inducible degradation (AID) sequence at the N-terminus of the IRS2 gene provided rapid and reversible control of IRS2 protein degradation to investigate the impact of inducible reduction of IRS2 expression on cellular function. Our studies demonstrate that acute loss of IRS2 expression reduces CSC numbers, confirming the role of IRS2 in the regulation of CSC function that was observed upon chronic knockout or knockdown of IRS2 (12). Knock-in of mNeon-green at the N-terminus facilitated the analysis of endogenous IRS2 expression and localization. Live fluorescence imaging of these cells revealed that IRS2 shuttles between the cytoplasm and nucleus, a function that had not been appreciated in previous studies. Specifically, IRS2 expression is primarily cytosolic when cells are grown in full growth medium, but it accumulates in the nucleus under growth deprivation conditions. Stimulation with IGF-1 decreases IRS2 expression in the nucleus and promotes a corresponding increase in cytosolic localization. A region in the IRS2 C-terminal tail that includes a predicted nuclear export signal is required for export of IRS2 from the nucleus, as IRS2 accumulates in the nucleus when this region is deleted. Together our data highlight the value of studying endogenously tagged IRS2 as a model to examine IRS2 regulation and function.

Our studies of endogenous IRS2 tagged with mNeon-green have revealed novel information about the regulation of IRS2 intracellular localization. In previous studies using fixed cells and indirect immunofluorescence, IRS2 expression was found to be primarily cytosolic with minimal nuclear localization (22). This localization pattern was also observed in FFPE sections of human breast tumors (9). Studying endogenous IRS2 expression in live cells without the introduction of potential artifacts from fixation or overexpression of exogenously tagged proteins has provided a more accurate assessment of the intracellular localization of this adaptor protein. In live cells imaged in complete growth medium, IRS2 localization was observed to be primarily cytosolic, as reported previously. However, analysis of cells after serum-starvation and subsequent stimulation with IGF-1 revealed a more dynamic regulation of IRS2 localization, with IRS2 nuclear accumulation correlating with growth deprivation conditions. These findings suggest that growth regulatory signals control the localization of IRS2, although the mechanism by which this occurs and the impact of this regulation on the signaling functions of IRS2 remain to be investigated. Interestingly, the dynamic regulation of IRS2 localization was more evident in MDA-MB-231 cells upon serum starvation than was observed in SUM-159 cells, which may reflect the different oncogenic backgrounds of these cell lines. Specifically, SUM-159 cells express an activating PIK3CA mutation which sustains PI3K pathway activity at higher baseline levels upon serum starvation (24). These mNeon-green knock-in cells will facilitate real-time imaging studies to examine regulatory mechanisms that control IRS2 localization. Analysis of the DFN cells *in vivo* will also allow for the investigation of IRS2 expression and localization at the cellular and whole tumor level.

The contributions of IRS2 to tumor cell biology have been revealed through studies using cell and mouse model systems in which the IRS2 gene has been genetically knocked out by Cre-lox recombination or CRISPR/Cas9-mediated gene editing (11,12,25). Through analysis of these models, a role for IRS2 in regulating breast cancer progression and metastasis has been revealed. IRS2 contributes to more aggressive tumor behavior through its ability to regulate CSC self-renewal to promote tumor initiation at both primary and metastatic sites and through the regulation of tumor cell invasion (11,12). However, the impact of acute loss of IRS2 function in established tumors, which would occur in the therapeutic setting, has not been assessed previously due to the absence of direct targeted inhibitors of this adaptor protein. By introducing an auxin-inducible degron into the IRS2 gene, we have developed a model system that allows for the inducible degradation and restoration of IRS2 expression in a temporal manner. We have demonstrated that acute degradation of IRS2 recapitulates the loss of CSC function that we previously observed with chronic IRS2 loss (12). These knock-in models that express IRS2 with both AID and mNeon-green tags will be useful both in the *in vitro* and *in vivo* setting to further explore IRS2 localization and function and its potential as a target for therapy.

## Methods and Materials

### Cells, antibodies and reagents

SUM-159 cells were obtained from BioIVT and cultured in F-12 medium (Gibco) supplemented with 10% FBS, HEPES (25 mM), insulin (5 µg/mL) and hydrocortisone (1 µg/mL). MDA-MB-231 cells were obtained from ATCC and cultured in RPMI medium (Gibco) supplemented with 10% FBS. Cells were authenticated by STR profiling at the University of Arizona Genetics Core. All cells were incubated at 37 °C in 5% CO_2_. Cells were screened for mycoplasma contamination using the Mycoplasma PCR Detection Kit (Applied Biological Materials, Inc.).

For auxin-induced degradation experiments, IAA Indole-3-acetic acid sodium (Sigma) was dissolved in NaOH (1 M) solution to a concentration of 500 mM and cells were incubated with 0.5 mM IAA prior to protein extraction. For washout experiments, cells were treated with 0.5 mM IAA for 6 h, washed with PBS and incubated further in complete culture medium before collection for immunoblot analysis.

Primary antibodies: rabbit anti-IRS2 (Cell Signaling Technology), rabbit anti-mNeonGreen (E8E3V) (Cell Signaling Technology), mouse anti-GAPDH (Santa cruz), mouse anti-pTyr (Santa cruz), rabbit anti-p85 (Cell Signaling Technology), rabbit anti-AKT (Cell Signaling Technology), rabbit anti-pS473 AKT (Cell Signaling Technology), mouse anti-FLAG (Sigma). Secondary antibodies: anti-rabbit IgG HRP (Jackson ImmunoResearch), anti-mouse IgG HRP (Jackson ImmunoResearch).

### Construction of gRNA and donor plasmid

gRNAs for IRS2 were designed using the CRISPR tool implemented in CRISPOR (26). The N-terminal genomic region of the IRS2 gene was targeted. The most efficient gRNA was selected that cleaves within 20 bp of the start codon (gRNA sequences can be found in Table 1, gRNA efficiency measured by TIDE analysis) (27). crRNA and tracrRNA were purchased from IDT and made duplex for gRNA. A donor plasmid containing the AID:3X-FLAG:mNeon-green:linker nucleotide sequences flanked by left and right homology regions representing 789 bp upstream and 844 bp downstream of the IRS2 start codon was generated by Azenta and used to target the N-terminal region of the IRS2 gene by Cas9-mediated HDR. The IRS2 start codon was removed from the genomic sequence of the right homology arm. A donor fragment was generated from the plasmid template by PCR using Powerpol 2X PCR mix (ABclonal) (primer sequences can be found in Table 1). The parameters were as follows: 98 °C for 45 s, then 30 cycles of 98 °C for 10 s, 62.8 °C for 30 s, 72°C for 90 s (30 s/kb), and a final step at 72°C for 5 min. For auxin receptor F-box protein expression, gRNA (19) and pSH-EFIRES-P-AtAFB2 plasmid were used (Addgene) (17).

**Table 1.**
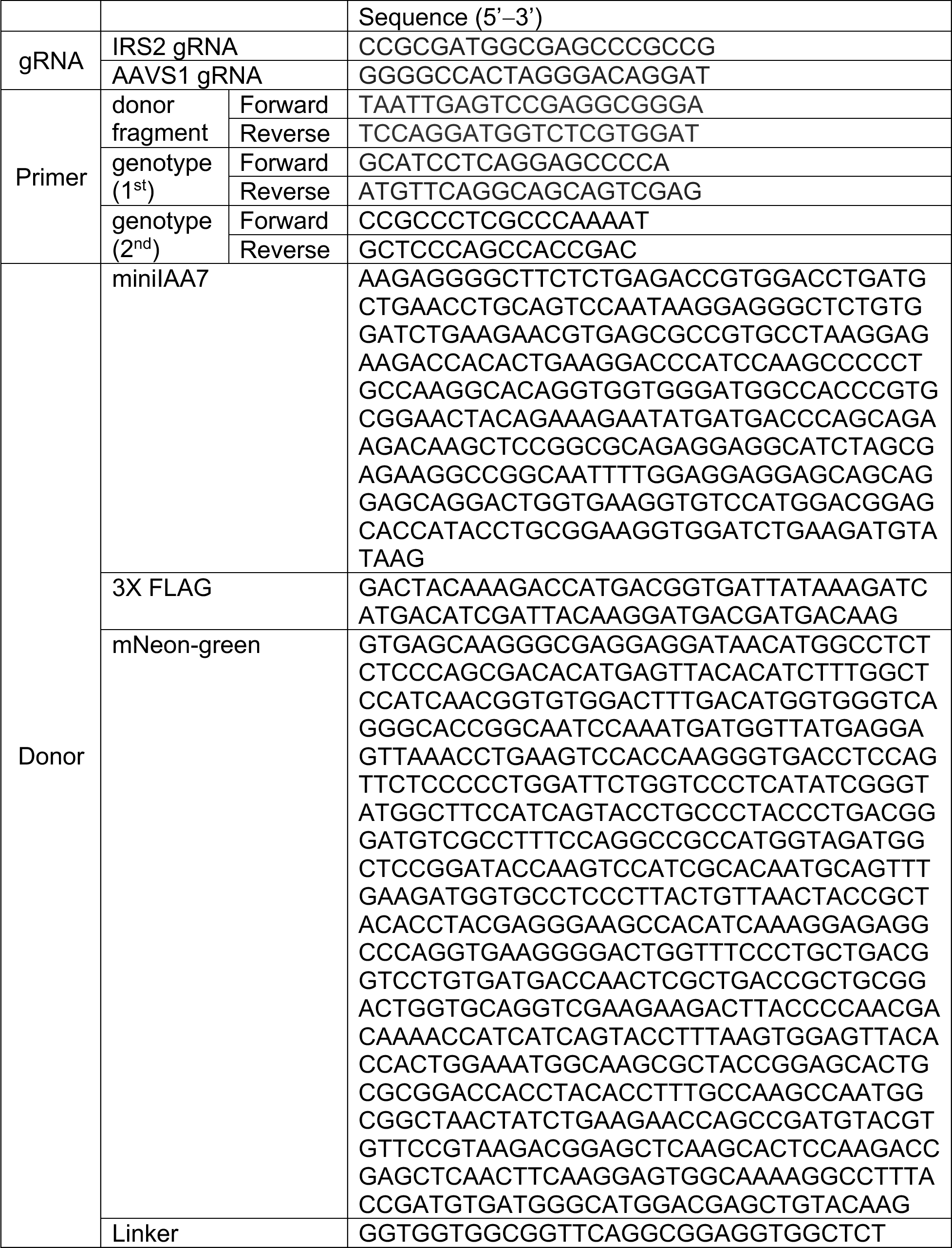
gRNA, Primer and Donor sequences.

### Generation of IRS2 knock-in cells

1.0 × 10^6^ cells were electroporated with 240 pmol crRNA: tracrRNA duplex,103.7 pmol S.p. Cas9 Nuclease V3 (IDT) and 3.0 μg donor fragment per 100 μL of Nucleofector™ Solution V (Lonza) using the Amaxa Nucleofector II (Lonza). For MDA-MB-231 cells, 60 pmol Cas9 Electroporation enhancer (IDT) was included. After electroporation, 500 µL of prewarmed culture media was added per cuvette and the cells were gently resuspended and transferred to a 6-well plate with 2 mL culture media (SUM-159) and 1.17 pmol HDR enhancer V2 (IDT) (MDA-MB-231). Cells were incubated for 24 hours (SUM-159) or 48 hrs (MDA-MB-231) prior to sorting mNeon-green positive cells into 96 well plates using the FACSMelody™ (BD Biosciences). Single cell clones were expanded and screened by PCR for homozygous knock-in of the DFN cassette.

Homozygously tagged clones were used to express the auxin receptor F-box protein AtAFB2 in the AAVS1 safe harbor locus. 1.0 × 10^6^ cells were electroporated as above with 3.0 μg AtAFB2 plasmid, gRNA and Cas9. Electroporated cells were plated into 10 cm plates and incubated for 24 hours prior to selection with 1 μg/mL puromycin.

### Genotyping PCR

Cells were lysed in QuickExtract™ DNA Extraction Soln 1.0 (BIOSEARCH Technologies) for 15 min at 68 °C and 15 min at 98 °C. Target regions were amplified by PCR using Powerpol 2X PCR mix. The parameters were as follows: 98 °C for 45 s, then 30 cycles of 98 °C for 10 s, 71 °C for 30 s, 72 °C for 1 min (30 s/kb), and a final step at 72 °C for 5 min for focused on tag region (1^st^ round PCR) and 98°C for 45 s, then 35 cycles of 98 °C for 10 s, 66°C for 30 s, 72°C for 2 min (30 s/kb), and a final step at 72°C for 5 min for over insert region (2^nd^ round PCR). The amplified products were analyzed by agarose gel electrophoresis.

### Immunoblotting

Cells were seeded at 1 × 10^5^ cells/well in a 6-well plate and grown for 24 hrs before protein extraction in lysis buffer (20 mM Tris-HCl pH 7.4, 137 mM NaCl, 10% Glycerol, and 1% NP-40) containing 1x complete Mini (Roche) and 1x PhosSTOP (Roche). Lysates were centrifuged at 13,000 rpm for 10 min at 4°C and the protein concentrations were quantified using Pierce BCA Protein Assay Kit (Thermo Fisher). Equal amounts of protein were separated on 8% SDS-PAGE and then transferred onto nitrocellulose membranes (Thermo Fisher). After blocking in 5% milk/TBST at room temperature for 1 h, the membrane was incubated with primary antibodies at 4°C overnight and subsequently incubated with HRP-conjugated secondary antibodies at room temperature for 1 h and all images were acquired using a ChemiDoc XRS+ system (Bio-Rad). Mouse anti-GAPDH antibody was used at a 1:2000 dilution with TBST containing 3% BSA (Sigma), and other primary antibodies were used at a 1:1000 dilution. All secondary antibodies were used at a 1:5000 dilution with TBST containing 3% BSA.

### Fluorescence microscopy

For live cell imaging, cells plated in glass bottom plates were incubated overnight in either complete culture medium or medium lacking FBS (serum starved) and then viewed by confocal microscopy (Nikon ECLIPSE Ti2; 60X). Cells in serum-free medium were stimulated with IGF-1 and imaged at set time points (5-30 min). For fixed cell imaging, cells in 8-well chamber glass slides were washed three times with Dulbecco’s PBS and fixed in 3.8% paraformaldehye in Dulbecco’s PBS with 0.5% Tween (PBST) for 1 hr. Permeabilized cells were blocked for 1 hr using 3% BSA in PBST and then incubated with anti-HA antibodies diluted in blocking buffer at room temperature for 1 hr. Secondary antibodies were diluted in the same buffer and cells were incubated at room temperature for an additional 30 minutes. Cells were washed three times with PBST after each antibody incubation. Coverslips were then mounted onto the glass slides using Prolong Gold containing DAPI (Cell signaling) and the slides were viewed by confocal microscopy. All images were adjusted equally for brightness/contrast using ImageJ software. The expression intensity of IRS2 in the cytoplasm and nucleus was measured using ImageJ software.

### Limiting dilution assay

Single-cell suspensions were plated in ultra-low-attachment 96-2 well plates (Corning) in serial dilutions (5, 10, 25, 50, 100 cells/well) in B27 mammosphere media. After 5 days, wells were imaged using a Celigo imaging cytometer and the number of wells containing ≥1 mammosphere were determined. Cancer stem cell frequency was determined using ELDA (28).

### Data analysis

Statistical analysis was performed using Prism10.0, Graphpad. Two-way ANOVA with Dunnett’s post-test were applied and p-value of <0.05 was considered to indicate statistical significance.

## Acknowledgements

We thank Art Mercurio and the Mercurio lab for helpful discussion and comments on the manuscript. This work was supported by National Institute of Health (NIH) grants CA227993 and CA229910 (LMS). The content is solely the responsibility of the authors and does not necessarily represent the official views of the National Institutes of Health.

## Author Contributions

MJ and LMS were involved in the conception and design of the project and wrote the manuscript; J-SL, MWL, and JSM were involved with the development of methodology and the acquisition and analysis of data. All authors reviewed and approved the final version of the manuscript.

---

**Supplemental Figure 1. Generation and characterization of DFN-KI cells.** A. TIDE analysis for the gRNA used to knock-in the DFN cassette by homology directed repair. B. Density plots for the sorting of mNeon-green positive cells. C. Flow cytometry analysis for mNeon green in WT-IRS2 cells, DFN-IRS2 cells and DFN-IRS2 cells after a 6 hrs treatment with auxin.

## References

1. Lee, J. S., Tocheny, C. E., and Shaw, L. M. (2022) The Insulin-like Growth Factor Signaling Pathway in Breast Cancer: An Elusive Therapeutic Target. Life (Basel) 12

2. Lero, M. W., and Shaw, L. M. (2021) Diversity of insulin and IGF signaling in breast cancer: Implications for therapy. Mol Cell Endocrinol 527, 111213

3. Schnarr, B., Strunz, K., Ohsam, J., Benner, A., Wacker, J., and Mayer, D. (2000) Down-regulation of insulin-like growth factor-I receptor and insulin receptor substrate-1 expression in advanced human breast cancer. Int J Cancer 89, 506–513

4. Sisci, D., Morelli, C., Garofalo, C., Romeo, F., Morabito, L., Casaburi, F., Middea, E., Cascio, S., Brunelli, E., Ando, S., and Surmacz, E. (2007) Expression of nuclear insulin receptor substrate 1 in breast cancer. J Clin Pathol 60, 633–641

5. Byron, S. A., Horwitz, K. B., Richer, J. K., Lange, C. A., Zhang, X., and Yee, D. (2006) Insulin receptor substrates mediate distinct biological responses to insulin-like growth factor receptor activation in breast cancer cells. Br J Cancer 95, 1220–1228

6. Wu, A., Chen, J., and Baserga, R. (2007) Nuclear insulin receptor substrate-1 activates promoters of cell cycle progression genes. Oncogene

7. Cesarone, G., Garofalo, C., Abrams, M. T., Igoucheva, O., Alexeev, V., Yoon, K., Surmacz, E., and Wickstrom, E. (2006) RNAi-mediated silencing of insulin receptor substrate 1 (IRS-1) enhances tamoxifen-induced cell death in MCF-7 breast cancer cells. J Cell Biochem 98, 440–450

8. Porter, H. A., Perry, A., Kingsley, C., Tran, N. L., and Keegan, A. D. (2013) IRS1 is highly expressed in localized breast tumors and regulates the sensitivity of breast cancer cells to chemotherapy, while IRS2 is highly expressed in invasive breast tumors. Cancer Lett 338, 239–248

9. Clark, J. L., Dresser, K., Hsieh, C. C., Sabel, M., Kleer, C. G., Khan, A., and Shaw, L. M. (2011) Membrane localization of insulin receptor substrate-2 (IRS-2) is associated with decreased overall survival in breast cancer. Breast Cancer Res Treat 130, 759–772

10. Nagle, J. A., Ma, Z., Byrne, M. A., White, M. F., and Shaw, L. M. (2004) Involvement of insulin receptor substrate 2 in mammary tumor metastasis. Mol Cell Biol 24, 9726–9735

11. Mercado-Matos, J., Janusis, J., Zhu, S., Chen, S. S., and Shaw, L. M. (2018) Identification of a novel invasion-promoting region in Insulin Receptor Substrate 2 (IRS2). Mol Cell Biol

12. Lee, J. S., Lero, M. W., Mercado-Matos, J., Zhu, S., Jo, M., Tocheny, C. E., Morgan, J. S., and Shaw, L. M. (2022) The insulin and IGF signaling pathway sustains breast cancer stem cells by IRS2/PI3K-mediated regulation of MYC. Cell Rep 41, 111759

13. Zhu, S., Ward, B. M., Yu, J., Matthew-Onabanjo, A. N., Janusis, J., Hsieh, C. C., Tomaszewicz, K., Hutchinson, L., Zhu, L. J., Kandil, D., and Shaw, L. M. (2018) IRS2 mutations linked to invasion in pleomorphic invasive lobular carcinoma. JCI Insight 3

14. Ma, Z., Gibson, S. L., Byrne, M. A., Zhang, J., White, M. F., and Shaw, L. M. (2006) Suppression of insulin receptor substrate 1 (IRS-1) promotes mammary tumor metastasis. Mol Cell Biol 26, 9338–9351

15. Bukhari, H., and Muller, T. (2019) Endogenous Fluorescence Tagging by CRISPR. Trends Cell Biol 29, 912–928

16. Nishimura, K., Fukagawa, T., Takisawa, H., Kakimoto, T., and Kanemaki, M. (2009) An auxin-based degron system for the rapid depletion of proteins in nonplant cells. Nat Methods 6, 917–922

17. Li, S., Prasanna, X., Salo, V. T., Vattulainen, I., and Ikonen, E. (2019) An efficient auxin-inducible degron system with low basal degradation in human cells. Nat Methods 16, 866–869

18. Yesbolatova, A., Saito, Y., Kitamoto, N., Makino-Itou, H., Ajima, R., Nakano, R., Nakaoka, H., Fukui, K., Gamo, K., Tominari, Y., Takeuchi, H., Saga, Y., Hayashi, K. I., and Kanemaki, M. T. (2020) The auxin-inducible degron 2 technology provides sharp degradation control in yeast, mammalian cells, and mice. Nat Commun 11, 5701

19. Yunusova, A., Smirnov, A., Shnaider, T., Lukyanchikova, V., Afonnikova, S., and Battulin, N. (2021) Evaluation of the OsTIR1 and AtAFB2 AID Systems for Genome Architectural Protein Degradation in Mammalian Cells. Front Mol Biosci 8, 757394

20. Sawka-Verhelle, D., Tartare-Deckert, S., White, M. F., and Van Obberghen, E. (1996) Insulin receptor substrate-2 binds to the insulin receptor through its phosphotyrosine-binding domain and through a newly identified domain comprising amino acids 591-786. J Biol Chem 271, 5980–5983

21. Mardilovich, K., Pankratz, S. L., and Shaw, L. M. (2009) Expression and function of the insulin receptor substrate proteins in cancer. Cell Commun Signal 7, 14

22. Mercado-Matos, J., Clark, J. L., Piper, A. J., Janusis, J., and Shaw, L. M. (2017) Differential involvement of the microtubule cytoskeleton in insulin receptor substrate 1 (IRS-1) and IRS-2 signaling to AKT determines the response to microtubule disruption in breast carcinoma cells. J Biol Chem 292, 7806–7816

23. Xu, D., Marquis, K., Pei, J., Fu, S. C., Cagatay, T., Grishin, N. V., and Chook, Y. M. (2015) LocNES: a computational tool for locating classical NESs in CRM1 cargo proteins. Bioinformatics 31, 1357–1365

24. Lehmann, B. D., Bauer, J. A., Chen, X., Sanders, M. E., Chakravarthy, A. B., Shyr, Y., and Pietenpol, J. A. (2011) Identification of human triple-negative breast cancer subtypes and preclinical models for selection of targeted therapies. J Clin Invest 121, 2750–2767

25. Landis, J., and Shaw, L. M. (2014) Insulin receptor substrate 2-mediated phosphatidylinositol 3-kinase signaling selectively inhibits glycogen synthase kinase 3beta to regulate aerobic glycolysis. J Biol Chem 289, 18603–18613

26. Concordet, J. P., and Haeussler, M. (2018) CRISPOR: intuitive guide selection for CRISPR/Cas9 genome editing experiments and screens. Nucleic Acids Res 46, W242–W245

27. Brinkman, E. K., and van Steensel, B. (2019) Rapid Quantitative Evaluation of CRISPR Genome Editing by TIDE and TIDER. Methods Mol Biol 1961, 29–44

28. Hu, Y., and Smyth, G. K. (2009) ELDA: extreme limiting dilution analysis for comparing depleted and enriched populations in stem cell and other assays. J Immunol Methods 347, 70-78

