## Supplemental Figure 1 for "Fluorescent tagging of endogenous IRS2 with an auxin-dependent degron to assess dynamic intracellular localization and function"

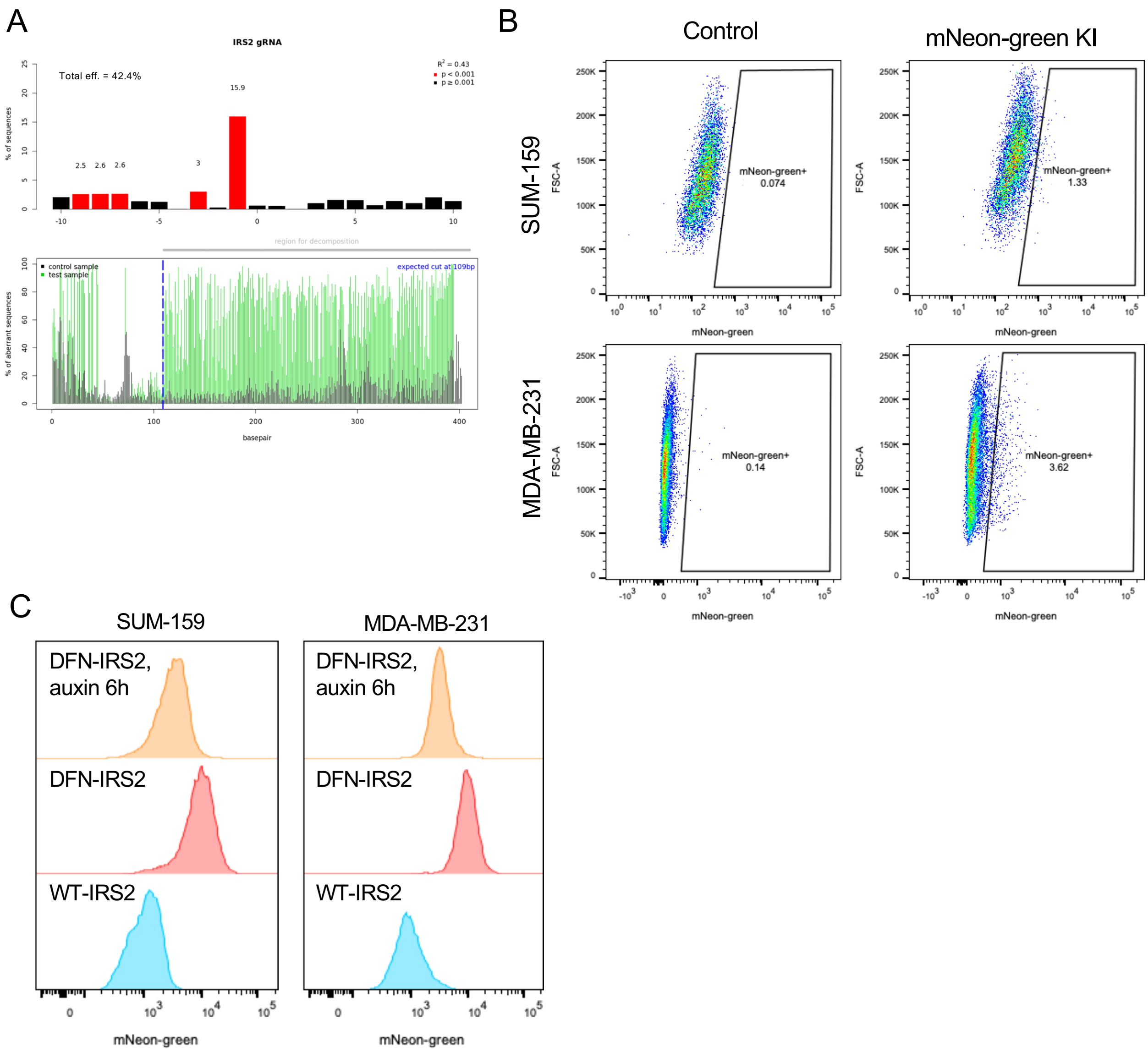


**Supplemental Figure 1. Generation and characterization of DFN-KI cells.** A. TIDE analysis for the gRNA used to knock-in the DFN cassette by homology directed repair. B. Density plots for the sorting of mNeon-green positive cells. C. Flow cytometry analysis for mNeon green in WT-IRS2 cells, DFN-IRS2 cells and DFN-IRS2 cells after a 6 hrs treatment with auxin.
